# Osmium Tetroxide-Enhanced µCT Enables 3D Mapping of Intra-articular Adipose Tissue

**DOI:** 10.64898/2026.11.24.768417

**Authors:** Danny M. Nguyen, Michael D. Newton, Madison Buckles, Isabelle J. Smith, Katherine Toufanian, Demetria Tonelli, Alexander J. Knights, Tristan Maerz

## Abstract

**Objectives:** Intra-articular adipose tissue is a unique adipose depot that has received increasing attention in the context of osteoarthritis pathophysiology, but no robust methodologies currently exist for its 3D visualization and quantification in preclinical models. This study developed a protocol for visualization and quantification of intra-articular adipose tissue in murine knee joints using osmium tetroxide (OsTet)-based contrast-enhanced micro-computed tomography (µCT), followed by illustration of its utility to assess adipose remodeling in a murine post-traumatic osteoarthritis (PTOA) model.

**Design:** Systematic optimization of sample preparation, OsTet incubation conditions, and µCT acquisition parameters was conducted to establish a protocol tailored for murine knee joints. PTOA was induced via non-invasive anterior cruciate ligament rupture to demonstrate the utility of this protocol in the assessment of disease-associated remodeling.

**Results:** Optimization of our OsTet-enhanced µCT protocol overcame attenuation artifacts and incomplete penetrance in intra-articular adipose depots of the murine knee. Adjusting acquisition parameters (0.10 mm Cu filter, 90 kVp) mitigated beam hardening and resolved tissue heterogeneity. Complete tissue penetration and uptake equilibrium were achieved using a 1% OsTet concentration incubated for 48 h in both healthy and injured limbs. Applied to a PTOA model, the pipeline reliably detected microstructural adipose remodeling, with low signal regions corresponding to fibrotic tissue, which was validated by histopathological staining.

**Conclusions:** This study established OsTet-enhanced µCT as a reliable imaging method for joint-wide assessment of adipose remodeling, providing a valuable framework for future studies investigating the role of joint adipose tissue in development, homeostasis, and disease.

## Introduction

Remodeling of adipocyte-rich tissue is associated with various inflammatory and fibrotic conditions^1,2^, and in the context of joint diseases, the homeostatic and pathogenic roles of adipose tissue are garnering attention. In conditions such as osteoarthritis (OA), both systemic and local adipose tissue are increasingly recognized as regulators of disease onset and progression^3^. Local adipocyte-derived signaling was shown to maintain homeostatic fibroblast programs in healthy synovium, whereas OA promotes changes in adiposity, altered adipocyte and adipose-resident stromal cell states, and disrupted adipose metabolism^4^. Parsing the effects of local intra-articular adipose tissue in OA from systemic depots has prompted the development of novel preclinical models and next-generation sequencing datasets, further elucidating the unique cellular and molecular interactions in homeostatic and pathologic adipose within the joint microenvironment^5,6^.An association between adipose tissue dysfunction and OA pain is also emerging (reviewed in^7^). Together these findings support the growing notion that remodeling of intra-articular adipose tissue contributes to the multi-faceted pathologies associated with OA onset and progression, emphasizing the importance of further studies focused on intra-articular adipose remodeling.

MRI and ultrasound sonography have been used in preclinical and clinical research to characterize structural changes of intra-articular adipose tissue throughout OA progression – most notably, focusing on the infrapatellar fat pad (IFP), the most prominent and best studied intra-articular adipose depot. However, these modalities primarily capture gross tissue-level changes and fail to resolve more specific aspects of the adipose phenotype. MRI studies have reported overall IFP volume loss in preclinical gout^8^, and clinical studies have demonstrated that both IFP volume and contrast uptake from dynamic contrast-enhanced MRI positively correlate with OA-associated pain^9^. Similarly, ultrasound elastography has shown that increased IFP stiffness is associated with pain severity in OA^10^. Despite these insights, both modalities have important limitations. Conventional MRI signal reflects both lipid and water content, and in inflamed joints, edema contributes substantially to signal intensity, limiting specificity for adipose tissue. Ultrasound elastography likewise provides an indirect, operator-dependent measure of bulk tissue stiffness that cannot distinguish adipose loss from fibrosis, edema, or other compositional changes within the IFP. Thus, these modalities lack specificity in the direct assessment of adipose tissue, and in the setting of preclinical studies, they also lack the necessary spatial resolution for small tissue volume, highlighting the need for alternative imaging strategies.

Histology remains the gold standard to derive detailed tissue-level information about adipose tissue in preclinical models; however, it is inherently limited to two-dimensional evaluation, precluding an analysis of joint-wide adipose changes. Furthermore, while most preclinical and clinical studies have focused on the IFP, there are multiple adipose depots (colloquially referred to as ‘fat pads’) in the knee joint, including the posterior fat pad (PFP), suprapatellar fat pad, and pericruciate fat pads. Volumetric and compositional changes to these depots are understudied and more challenging to holistically capture in 2D histological sections due to their diffuse distribution. These constraints highlight the need for a high resolution, three-dimensional (3D) imaging approach that can derive disease-relevant quantitative information from intra-articular adipose tissue in preclinical models.

Osmium tetroxide (OsTet) is a heavy metal, lipophilic molecule that preferentially binds to unsaturated fatty acids, leading to the reduction of osmium esters and deposition of a dense, black precipitate^11^. Given its high electron density, OsTet exhibits high radiodensity, enabling clear distinction of lipid-rich tissues from surrounding structures on radiographic imaging. Originally introduced by George Palade in 1952 for histologically visualizing cell membranes^12^, OsTet has since been widely adopted for visualization of adipose tissue and lipids, including in myelin sheaths for peripheral nerve anatomy studies^13^ and bone marrow adipose tissue quantification in murine long bones^14^, demonstrating its utility across multiple research domains.

Despite growing evidence that intra-articular adipose tissue contributes to joint homeostasis and disease, compartment-specific 3D remodeling of these tissues during arthritis progression remains poorly characterized. To date, OsTet-enhanced µCT has not been used to morphometrically assess intra-articular adipose depots. To this end, the objective of this study was to establish an OsTet-enhanced, high-resolution µCT workflow for 3D visualization and quantification of intra-articular adipose tissue in the murine knee joint.

## Method

### Animal Usage

All work described in this study was conducted under a protocol approved by the University of Michigan IACUC. Mice were housed in a 12 h dark/light facility and allowed *ad libitum* activity, food, and water. 12 to 20-week-old, functionally wild-type male mice (C57BL/6 background) were used for each OsTet optimization study. 12-week-old, male C57BL/6 mice were used for the PTOA model study, and contralateral limbs were used as internal controls (n=6 ACLR joints, n=6 contralateral joints).

### Hindlimb Harvest and Dissection

Following euthanasia by CO_2_ asphyxiation, whole hindlimbs were obtained by disarticulating the hip joint, and skin was removed. Preliminary analyses demonstrated incomplete penetrance of OsTet into the joint, particularly in the posterior aspect of the joint, if excessive muscle was retained (**Fig. S1A, B**). Therefore, extra-articular muscle and connective tissue were removed by severing all tendons located in the ankle and individually peeling back the tibialis anterior, lateral/medial gastrocnemius, and soleus along the tibia and cutting at their respective peri-articular insertions (**Fig. S1C, D**). Approximately 80% of the quadriceps was removed, leaving the most distal part of the quadriceps and the quadriceps tendon intact to avoid compromising the synovial capsule (**Fig. S1D**). Samples were then fixed in 10% neutral-buffered formalin for 24 h. Subsequent OsTet staining and µCT of these more extensively dissected samples confirmed successful OsTet penetrance while preserving joint structure (**Fig. S1E**).

### OsTet Staining

Fixed limbs were either stored in 70% ethanol for storage or directly transferred to the OsTet staining solution. Following optimization, the final OsTet staining protocol was as follows:

1. Prepare OsTet incubation tubes in 50 mL conical tubes (20 mL total volume): 1% OsTet: 5 mL of 4% stock OsTet + 15 mL Sorenson’s Phosphate Buffer Solution (SPBS).
2. Transfer joints into respective tubes of OsTet/SPBS, seal and incubate for 48 h.
3. Wash:

a. After 48 h, tip out OsTet into oil waste, using a disposable strainer to catch the joint.
b. Use designated OsTet forceps or disposable forceps to transfer joint into a 15 mL conical tube containing 10 mL SPBS.
c. Mix 2-3 times to wash superficial OsTet off the joint then dispose of buffer in corn oil/OsTet waste.
d. Add joint into fresh 15 mL tube containing 10 mL SPBS, rotate for 2 h at room temperature.
e. Tip out buffer (catch joint) then transfer joint into new 15 mL tube of 10 mL SPBS, rotate for 2 h at room temperature.
f. Tip off buffer (catch joint) then transfer joint into new 15 mL tube of 10 mL SPBS, sit overnight at room temperature.
4. Remove joint from overnight wash and store in 5 mL Eppendorf tube filled with 4 mL SPBS for long-term storage at 4°C.

All processing and incubation steps must be carried out within a chemical fume hood.

### μCT Image Acquisition

Hindlimbs were packed in 3D printed cassettes and submerged in SPBS, and μCT imaging was performed using a Bruker 1176 μCT imaging system. During protocol refinement, three source voltages were used during acquisition (see **Supplemental Methods**) with a 9 μm voxel resolution, 0.30-degree rotation step, and 3 frame averages. Final datasets were acquired at a 9 μm voxel resolution, 0.25-degree rotation step, 4 frame averages using the 0.10 mm Cu filter with 90 kVp source voltage, 278 μA source current, and 1850 ms exposure.

### 3D Image Segmentation/Analysis

All segmentations and quantifications were performed in Dragonfly 3D World (version 2024.1 Build 1627). Datasets were visualized using multi-planar reconstruction in axial, sagittal, and coronal viewing planes. Manual outlines were contoured on 2D axial images to divide the anterior and posterior region of the articular space. Superior and inferior boundaries were established at the physis of each long bone using the 2D sagittal view. A 20,000+ Arbitrary Units (AU) threshold was empirically determined to allow for accurate segmentation of contrast-enhanced adipose tissue while minimizing false detection of structures outside the region of interest (e.g. bone, soft tissue, background, etc.) and was used for all scans. Manual outlines and contrast-enhanced tissue thresholds were then intersected, and volume and attenuation data were exported. Representative images were generated from a 3D rendering, with extra-articular tissue removed to visualize only intra-articular structures. 2D views were generated using a grayscale gradient range of 0 - 21,500 Hounsfield Units (HU) to enhance contrast between bone and background. Attenuation values were converted from AU to HU using calibration scans of pure water and air.

### Optimization of OsTet Staining for Murine Knee Joints

To characterize how acquisition parameters affect the attenuation of OsTet-stained tissue, the knee joints of functionally wild-type mice aged 12-20 weeks (n=2/concentration) were dissected, stained in 0.01% or 0.05% OsTet and scanned with 0.50 mm Al, 1.00 mm Al, and 0.10 mm Cu filters (see **Supplemental Methods**). Average attenuation of OsTet+ tissue was measured as described above. In concentration series experiments, bilateral knee joints (n=2/concentration) were harvested, stained in 0.25%, 0.43%, 0.60%, 0.71%, 1.0% OsTet, and scanned. Volumetric data were separated into anterior/posterior regions, and attenuation data were reported as an average of total stained intra-articular fat (IFP + PFP). For time series experiments, unilateral knee joints (n=3) were harvested, incubated in staining solution, and scanned at multiple timepoints (2, 4, 8, 16, 24, 48 h). To minimize time out of staining solution, lower resolution scans were obtained for this step (18 μm, 0.50-degree rotation step, 3 frame averages). Masks were contoured over the anterior joint space, and within each mask, both the mean attenuation of the segmented foreground (20,000 AU+) and the total mean attenuation across the entire masked region were quantified. Equilibration was modeled from time-dependent attenuation data using a one-phase association equation:

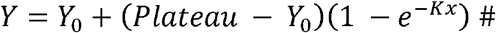

where *Y* represents experimentally measured attenuation, *Y*_0_ represents the initial attenuation at *X*=0 (constrained to 0), *Plateau* represents the asymptotic maximum attenuation, *K* represents the association rate constant (h^-1^), and *x* represents incubation time (h).

### Anterior Cruciate Ligament Rupture Model of Post-Traumatic Osteoarthritis

To induce PTOA, an established model of non-invasive anterior cruciate ligament rupture (ACLR) was utilized^15^. Mice were anesthetized with 2% isoflurane and subjected to unilateral ACLR via tibial compression overload in the right hindlimb. A single subcutaneous injection of carprofen (5 mg/kg, 1 mg/mL stock) was administered for analgesia immediately following the procedure. Contralateral limbs (left) were used as internal controls, as they demonstrate no detectable changes in PTOA severity or synovitis compared to sham mice^16^.

### Decalcification, Paraffin Processing and Histology

Following μCT imaging and quality control, limbs were decalcified in 100% Immunocal (Fisher Scientific, Cat. #: 50-980-229) for 12 h. Limbs were then sent to the University of Michigan Histology core for paraffin processing. Paraffin embedding and sagittal sectioning at a 5 μm thickness were performed in-house. Sections were collected from the weight bearing zone of the medial condyle/plateau, as well as from the trochlear groove (where the IFP is most prominent). Sections were stained using a Masson’s Trichrome protocol (EMS Catalog #: 26367-Series protocol). The protocol was modified by removing the scarlet acid fuchsin step as inclusion of that step demonstrated washout of OsTet. Slides were bright-field-imaged using a Keyence BZ-X800 microscope using a 10x objective, and images were stitched to allow for joint-wide visualization. Slides were qualitatively assessed for complete OsTet penetration and collagen deposition.

### Line Profile Analyses of Co-registered Images

To evaluate intra-articular tissue contrast retention and boundary definition, a semi-automated image processing pipeline was implemented in MATLAB (v. R2024b). Axial plane μCT slices were extracted from raw TIFF image stacks. To account for anatomical orientation, the contralateral joint served as the reference frame, while the ipsilateral joint slice was horizontally mirrored. To eliminate slice-selection bias and ensure identical anatomical sampling across joints, automated slice matching was performed using a structural gradient cross-correlation algorithm within a 30-slice search window around the target index of the injured joint. Matched slices were manually oriented to align the joint space vertically. Geometric registration between contralateral and ACLR slices was established via landmark control points (**Fig. S1G**) using a 2D similarity transformation. A line profile was drawn on the contralateral control image from the medial to lateral aspect of the anterior joint space and automatically transferred to the transformed ACLR slice coordinates. Intensity profiles were extracted along the vector using a 5-line parallel band-averaging filter (±2 pixel range, 1 pixel step) to eliminate single-pixel µCT reconstruction artifacts and maximize spatial signal-to-noise ratio. Quantitative outputs included attenuation plots along the drawn vector.

### Statistical analysis

All statistical analyses were conducted using GraphPad Prism 10.4.1. Unpaired, two-tailed parametric t-tests were performed to assess attenuation between intra-articular regions of interest, and all graphed results are reported as mean ± standard deviation (error bars). For regression curves, 95% confidence intervals are shown by minor dashed lines. One-way ANOVA with Tukey’s post-hoc comparison was used to compare attenuation values across varying acquisition groups. All comparisons met the assumptions of normality and homogeneity of variances. Statistical significance was defined as *P*<0.05, and exact *P*-values are provided for all significant comparisons.

Equilibration and regression analyses were performed in GraphPad Prism via a one-phase association model. Nonlinear curves were generated using least-squares regression and mean values, standard deviations, and 95% confidence intervals were plotted accordingly.

## Results

### Refinement of acquisition parameters and OsTet concentration

An OsTet-enhanced µCT protocol was previously established for three-dimensional imaging of bone marrow adipose tissue^14^ and has been widely adopted for visualization of lipid-rich structures. We therefore initially tested whether this existing protocol could be directly applied to intra-articular adipose tissue in the murine knee. Qualitative observations demonstrated incomplete penetrance of OsTet into all intra-articular regions coupled with excessive tomographic attenuation of contrast-enhanced tissue, resulting in detector saturation and prominent beam-hardening artifacts that obscured tissue architecture (**Fig. S1A**), underscoring the need to develop a knee joint-specific protocol and tailor the staining conditions, scanning parameters, and image analysis pipeline for this purpose (**Fig. 1A**).

**Figure 1.**
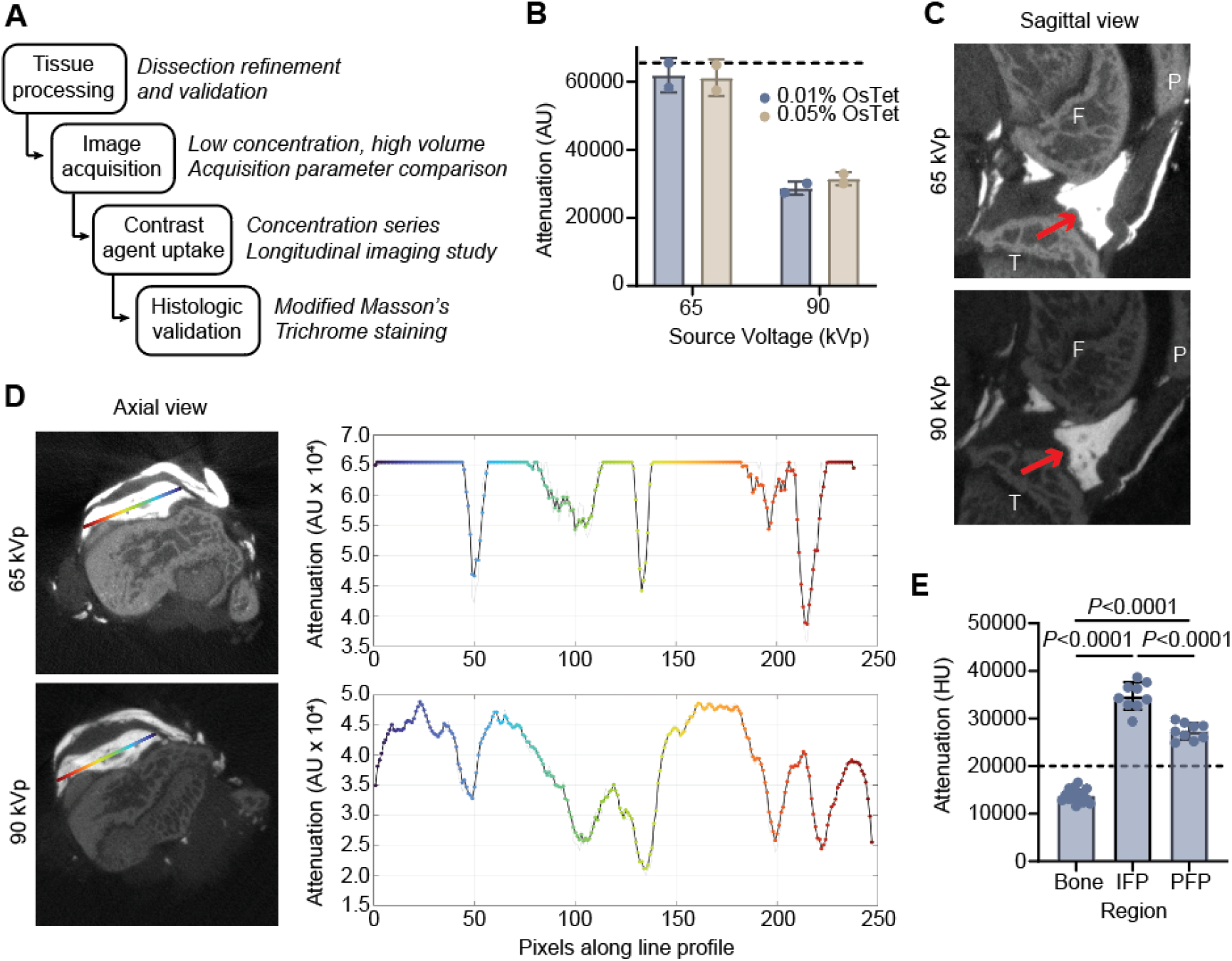
Refinement of μCT acquisition parameters. (**A**) Flow diagram of OsTet optimization phases. (**B**) Attenuation values from high volume, low concentration OsTet staining between 65 kVp and 90 kVp source voltage acquisitions. AU: arbitrary units. Dashed line represents the maximum attenuation threshold of the μCT scanner. (**C**) Representative sagittal μCT slices between 65 kVp and 90 kVp scans, with infrapatellar fat pad (IFP) marked by red arrow. P: patella, F: femur, T: tibia. (**D**) Left – Axial μCT slices with line profile overlays of 65 kVp and 90 kVp acquisitions. Right – Attenuation plots derived from line profiles. (**E**) Attenuation from three relevant regions of interest in the joint space (bone, IFP, and posterior fat pad (PFP)). Dashed line represents the minimum attenuation threshold used to define true fat signal.

To mitigate attenuation-related artifacts, we first tested markedly lower OsTet concentrations while maintaining high contrast agent volumes. Hindlimbs underwent OsTet staining at low concentrations (0.01% or 0.05%) in excess total volume (200 mL), followed by imaging. Unexpectedly, adipose tissue attenuation remained elevated with near-saturation signal observed even at 0.01% (**Fig. 1B**). This prompted us to mitigate overattenuation strictly through acquisition parameter adjustments. Comparative assessment of scans acquired with different source voltage and filter combinations demonstrated a significant reduction in adipose attenuation from ∼62,000 AU (1.00 mm Al filter, 65 kVp) to ∼30,000 AU (0.10 mm Cu filter, 90 kVp) (**Fig. 1B**). Resolution of attenuation-related imaging artifacts using these acquisition parameters resulted in imaging data that resolved intra-tissue heterogeneity and permitted granular delineation between true adipose tissue, connective tissue, and vasculature (**Fig. 1C**). Line profile analysis of slice-matched regions across heterogeneous contrast-enhanced tissue further demonstrated that acquisition at 65 kVp (Al filter) resulted in abundant detector saturation that obscured local signal differences, whereas 90 kVp (Cu filter) preserved attenuation profiles to improve signal contrast and detail at inter- and intra-tissue boundaries (**Fig. 1D**). The refined imaging protocol resulted in clear separation and significant differences between mean attenuation values of bone and contrast-enhanced adipose tissue in the anterior and posterior compartments (**Fig. 1E**).

### Optimization of OsTet concentration to facilitate whole-volume contrast uptake

We next sought to identify a suitable OsTet concentration that would result in repeatable enhancement of all intra-articular adipose regions by enabling complete OsTet penetration. To establish intra-versus extra-articular boundaries in 3D analysis, we defined superior and inferior bounds of the joint space as the femoral and tibial physes and the anterior and posterior bounds as the patellar tendon (anterior) and the edge of extra-articular soft tissue (posterior). Two masks were drawn, anterior and posterior, to segment the infrapatellar fat pad (IFP) and posterior fat pad (PFP) (pericruciate adipose tissue was not segmented separately in this analysis). A 20,000 AU threshold (corresponding to ∼7,000 Hounsfield units, HU) (**Fig. 1E**) was then applied within the drawn mask for segmentation and volumetric analyses (**Fig. 2A-B**).

**Figure 2.**
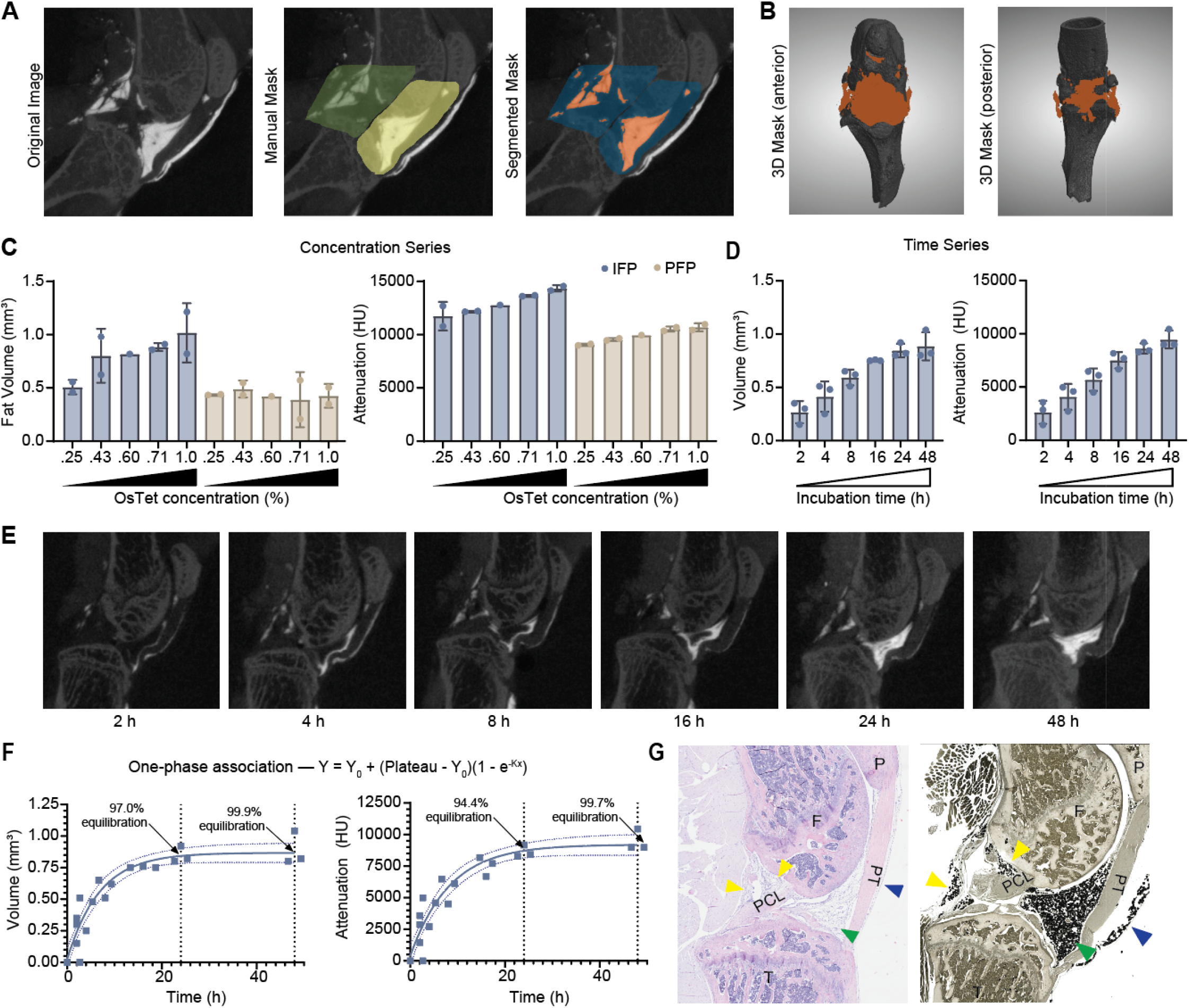
Optimization of OsTet concentration and incubation time. (**A**) Left to right: Representative sagittal μCT slices of original image, image with manual mask drawn (yellow: anterior manual mask; green: posterior manual mask), and segmented mask (orange: segmented foreground; blue: segmented background). (**B**) 3D renders of contrast-enhanced tissue post-segmentation, anterior (left) and posterior (right) aspect. (**C**) Contrast-enhanced positive fat signal volumes (left) and attenuation values (right) at increasing OsTet concentrations (24 h incubation, n=2 per concentration). (**D**) Contrast-enhanced positive fat signal volumes (left) and attenuation values (right) at increasing OsTet incubation times (1% OsTet concentration, n=3 per time point). (**E**) Representative sagittal μCT scans of OsTet-stained hindlimbs from 2 to 48 h (1% OsTet concentration). (**F**) Nonlinear regression models (one-phase association) fitted using volume and attenuation data (n=3). Equilibration extents are shown at 24 h and 48 h (large vertical dotted lines) and 95% confidence interval are shown (small dotted lines). (**G**) Representative sagittal histology sections depicting relevant intra-articular anatomy (H&E, left) alongside OsTet staining (right). P: patella, F: femur, T: tibia, PT: patellar tendon, PCL: posterior cruciate ligament. Green arrowhead: IFP, yellow arrowhead: PFP, blue arrowhead: subcutaneous prepatellar tendon fat.

Assessing OsTet concentrations across a 0.25%–1.0% range revealed a dose-dependent increase in contrast-enhanced IFP volume from 0.50 mm^3^ to 1.02 mm^3^, after which volumetric expansion reached a plateau, suggesting complete equilibration (**Fig. 2C**). The smaller, more diffuse PFP reached equilibration at ∼0.45mm^3^ at the lowest tested concentration (0.25%), indicating that the IFP should be the determining factor when selecting optimal OsTet concentration. With increasing concentration, IFP attenuation increased from ∼12,600 HU to ∼14,100 HU, and PFP attenuation increased from ∼9,000 HU to ∼10,500 HU, and attenuation profiles of all concentrations remained below the detector maximum of 25,400 HU (**Fig. 2C**). Analysis of imaging data and corresponding histological sections demonstrated that lower OsTet concentrations (0.25% and 0.43%) resulted in incomplete IFP staining, while the PFP was completely penetrated at all concentrations tested. Mid-range concentrations (0.60/0.71%) exhibited improved staining compared to lower concentrations; however, we noted distinct, circular pockets of unstained tissue in the center of the IFP (**Fig. S1F**). Staining voids were variable from sample to sample and were localized to the deepest regions of intra-articular adipose tissue, primarily within the IFP. Full tissue penetration was only observed at 1.0% OsTet, consistent with the observed increase in IFP volume up to 1.0% in the 3D data. Thus, we employed 1.0% OsTet concentration for all subsequent experiments.

### Longitudinal imaging identified contrast uptake equilibrium

When utilizing chemical contrast agents that rely on passive diffusion to generate tissue signal, high-quality imaging data is dependent upon the identification of contrast uptake equilibrium. Achieving equilibrium during contrast agent incubation minimizes imaging artifacts from ongoing diffusion during image acquisition, especially in lengthy acquisitions such as those in µCT imaging. To identify the timepoint at which OsTet uptake equilibrium is achieved in intra-articular adipose tissue, hindlimbs underwent longitudinal imaging at 2, 4, 8, 16, 24, and 48 h (**Fig. 2D, E**). At each incubation time point, samples were removed from OsTet solution, quickly rinsed, submerged and imaged in SPBS, and then returned to OsTet solution. This permitted matched, longitudinal analyses for rigorous determination of diffusion kinetics. Expectedly, contrast-enhanced tissue volume and attenuation of the IFP increased as time progressed, plateauing by 48 h (**Fig. 2D, E**). Using a one-phase association regression model, we observed that the IFP volume reached 97.0% of predicted equilibrium by 24 h and 99.9% by 48 h (R^2^ = 0.938) (**Fig. 2F, Table S1**). Modeling of attenuation data demonstrated similar behavior: high goodness-of-fit (R^2^ = 0.943) and equilibration predictions of 94.4% and 99.7% by 24 and 48 h, respectively (**Fig. 2F, Table S1**). Histologic analysis verified that complete staining with full tissue penetrance of OsTet throughout all intra-articular adipose depots was achieved by 48 h (**Fig. 2G**).

### OsTet-enhanced µCT identifies PTOA-associated fibrotic adipose remodeling

We next considered that fibrotic remodeling and increased extracellular matrix deposition, as seen in post-traumatic osteoarthritis (PTOA) and other joint diseases, could alter diffusion kinetics. We therefore performed further imaging in limbs from mice that underwent joint injury with our established non-invasive anterior cruciate ligament rupture (ACLR) model, which exhibits fibrotic remodeling of the synovium and IFP^17^. At 28 days (d) post-ACLR, a timepoint of established PTOA with extensive fibrotic remodeling of the IFP^18^, joints were processed and longitudinally imaged (**Fig. S2A**). Equilibration models based on normalized volume and attenuation indicated 99.6% (R^2^ = 0.7828) and 98.5% (R^2^ = 0.8893) equilibration, respectively, by 24 h, with volume and attenuation both reaching 99.9% equilibration by 48 h (**Fig. S2B, Table S2**), confirming that injured joints achieve complete contrast uptake equilibration within a 48 h incubation period, similar to uninjured joints. Histological staining confirmed complete penetrance of OsTet in ACLR samples, confirming that this contrast enhancement protocol is suitable for both PTOA and uninjured, contralateral limbs (**Fig. S2C**).

Further assessing the ability of OsTet-enhanced imaging to discern pathological IFP remodeling, line profile analysis between slice-matched contralateral and injured joints demonstrated marked, localized attenuation drop-offs throughout the IFP of 28d ACLR mice compared with contralateral limbs, corresponding to regions with non-adipocyte matrix deposition (**Fig. 3A**). Quantitatively, these signal drop-offs were large, with 30 to 60% reduction in attenuation, and some regions approached attenuation values of background, demonstrating that our OsTet imaging protocol resolves microstructural organization of joint adipose tissue, enabling nuanced 3D assessment of adipose remodeling.

**Figure 3.**
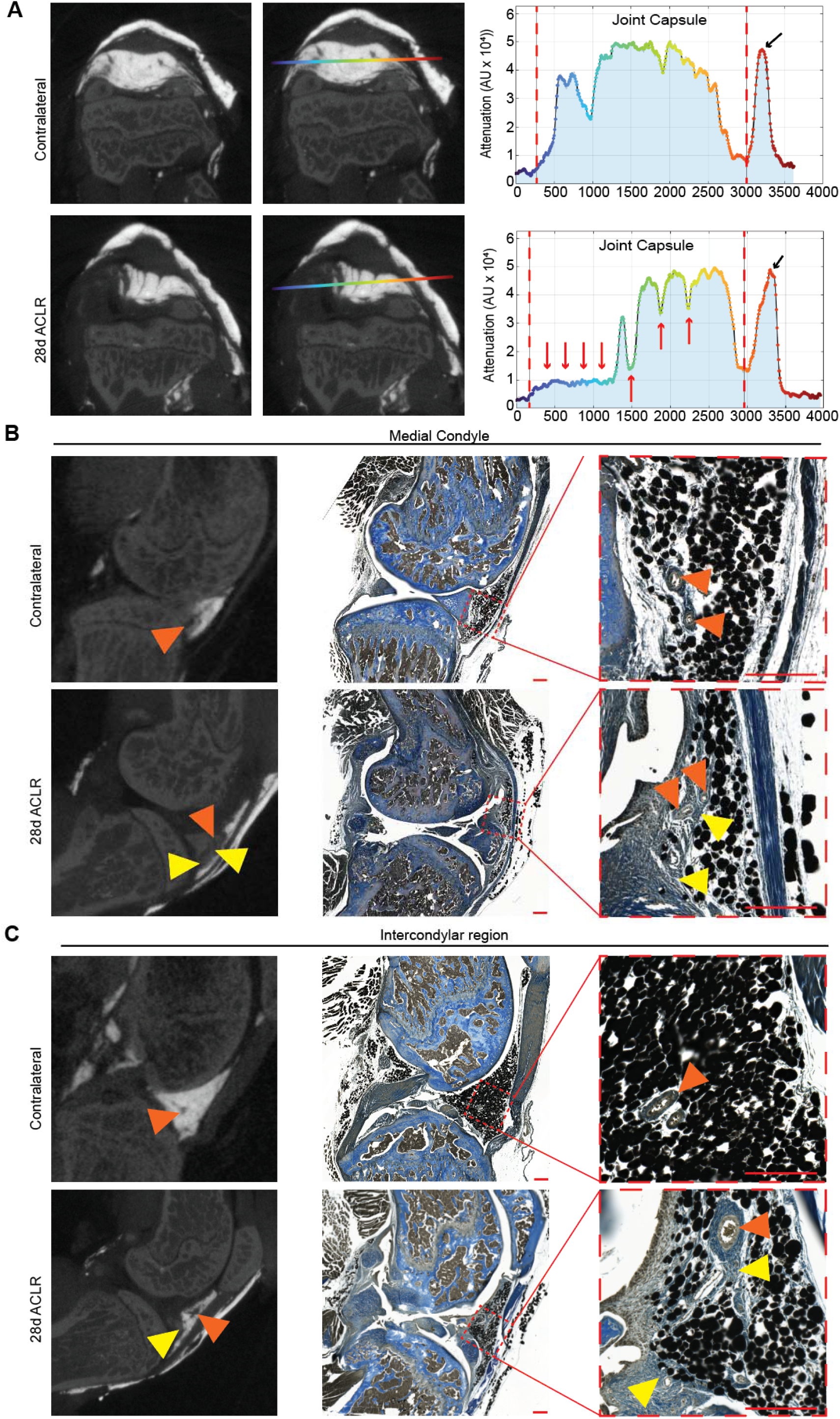
OsTet-enhanced µCT identifies PTOA-associated fibrotic adipose remodeling. (**A**) Representative axial μCT images of contralateral (top row) and 28d ACLR limbs (bottom row) with line profile overlay (middle) and attenuation profile outputs (right, AU: arbitrary units). Red arrows denote areas of attenuation drop off. Black arrows denote the subcutaneous (extra-articular) adipose tissue. Vertical dashed red lines represent joint capsule boundaries. (**B-C**) Left column: representative OsTet-stained sagittal μCT slices of contralateral and 28d ACLR from the medial condyle (**B**) or intercondylar region (**C**). Middle column: Slice-matched, modified Masson’s Trichrome and OsTet-stained histology sections. Scale bar: 100 μm. Right column: Zoomed in views of the IFP. Yellow arrowheads: absent OsTet staining in the IFP region co-localized with collagen deposition. Orange arrowheads: absent OsTet staining co-localized with vasculature in the IFP region.

Corroborating these imaging findings using structural histology, OsTet-stained sections were counterstained with a Modified Masson’s Trichrome and qualitatively evaluated for OsTet reduction and collagen deposition. In region- and depth-matched sections, the IFP of 28d ACLR joints exhibited a marked reduction of OsTet-positive adipocytes in both the medial and intercondylar joint compartments (**Fig. 3B-C**). In sections from contralateral uninjured joints, focal areas of low IFP attenuation corresponded to large blood vessels found in the healthy IFP. Expectedly, sections from 28d ACLR joints exhibited neovascularization and increased abundance of collagenous connective tissue infiltrating OsTet-stained adipose tissue, corresponding to regions of decreased imaging signal in matched µCT slices. These findings support the notion that fibrotic remodeling of intra-articular adipose tissue is detectable and quantitatively characterizable using our refined OsTet contrast enhancement and imaging protocol.

Taken together, our findings establish an optimized OsTet-enhanced µCT imaging workflow that provides robust and spatially resolved visualization of intra-articular adipose tissue, enabling 3D quantitative assessment of adipose tissue architecture and remodeling.

## Discussion

This study was motivated by the increased research focus on intra-articular adipose tissue and its role in joint homeostasis, adaptation, and disease. To enable more comprehensive and quantitative characterization of joint adipose tissue in murine models, we developed an osmium tetroxide-based μCT imaging workflow for three-dimensional visualization and quantification of intra-articular adipose tissue in the murine knee. Through systematic refinement of tissue preparation, imaging parameters, staining concentration, and incubation time, this work established a reproducible and sensitive method capable of resolving joint-wide adipose distribution and its disease-associated remodeling in a nondestructive manner.

Studies focused on various joint diseases, including OA, have implicated the IFP in joint homeostasis and disease pathogenesis^1,19,20^, but imaging-based analyses have been limited to 2D approaches. The methodological framework developed for this study is meant to contribute a valuable tool for the broader musculoskeletal research community to enable non-destructive assessment of intra-articular adipose in 3D prior to undertaking conventional histology. Historically, OsTet has been used to visualize lipid-rich structures in a 2D histological context, and more recently, for μCT imaging and 3D assessment of adipose tissue distribution and morphology in bone marrow and coronary artery structures^14,21^. This work expands OsTet’s utility in musculoskeletal research to reliably resolve intra-articular microarchitecture. Our protocol was able to capture regional tissue heterogeneity, especially in injured joints that exhibit fibrotic adipose remodeling, thereby offering a reproducible, non-destructive analysis of global adipose remodeling that cannot be reliably inferred from discrete histological sections alone. This approach can be readily adapted in other articular adipose depots such as the ankle, wrist, or elbow, or across other research domains for investigating lipid-rich tissues or disease models where spatial mapping of adipose remodeling is essential. While application of OsTet imaging of intra-articular joints could theoretically be extended to other species as well, the long incubation times required in this study suggest that reaching contrast uptake equilibrium may be challenging in larger animal models.

A minor modification of the conventional Masson’s Trichrome protocol enabled us to visualize fibrotic connective tissue alongside OsTet-stained adipocytes on histological sections, and other structural stains may be assessed in future studies. Thus, this protocol can likely be integrated into a standard histological processing pipeline during routine histopathological scoring. However, we did not assess whether immunostaining remains feasible. Several other limitations should be acknowledged. OsTet is a highly toxic and volatile substance which needs to be handled, stored, and disposed of with great care and with strict adherence to safety guidelines. Given its toxicity, its use is entirely limited to post-mortem *ex vivo* studies. As with all contrast-based imaging, contrast agent penetration and resultant signal intensity will vary with tissue composition, age, and disease state, necessitating careful refinement when adapting to new models. OsTet, as a lipophilic stain, binds independently of cell type, meaning that while lipid-rich adipocytes are its primary target, other lipid-laden cells may be stained, as well as extracellular free lipids. For this reason, it serves as a surrogate marker for adipocytes and adipose tissue, rather than a selective marker. Finally, while μCT provides high-resolution structural detail, it cannot directly capture the molecular or functional changes occurring within adipocytes or the surrounding stroma, highlighting the need to integrate complementary assays.

## Acknowledgments

We thank Dr. Kenneth Kozloff for use of the µCT imaging system and Dr. Erica Scheller for her guidance in our protocol refinement.

## Author Contributions

TM and AJK had full access to all the data included in this article and take full responsibility for the integrity of the data and accuracy of the information. All authors have been involved in the analysis and interpretation of the data and contributed to the final manuscript. Specific contributions to the manuscript are listed below, with author names given in the order in which they appear on the author line.

- Conception and design: DMN, MDN, AJK, TM.
- Analysis and interpretation of the data: DMN, MDN, AJK, TM.
- Drafting of the article: DMN, MDN, TM.
- Critical revision of the article for important intellectual content: All authors.
- Final approval of the article: All authors.
- Obtaining of funding: AJK, TM.

