## Supplementary Materials for "Osmium Tetroxide-Enhanced µCT Enables 3D Mapping of Intra-articular Adipose Tissue"

**Supplementary Methods**

**Supplementary Figures**

Figure S1

Figure S2

**Supplementary Tables**

Table S1

Table S2

**Supplementary Methods**

***μCT Parameters for Optimization Experiments***

0.50mm Al filter:

- Source voltage – 50 kVp
- Source current – 500 µA
- Exposure – 1350 ms

1.0mm Al filter:

- Source voltage – 65 kVp
- Source current – 385 µA
- Exposure – 1500 ms

0.10mm Cu filter:

- Source voltage – 90 kVp
- Source current – 278 µA
- Exposure – 1850 ms

Object to source distance was 120.451 mm and camera to source distance was 168.659 mm. Note: Voltage, current, and exposure were defaulted based on selected filter material and thickness. If default parameters are different, users can adjust each individual parameter using the scanner software.

***μCT Scan Reconstructions***

Reconstructions were performed using Bruker software, NRecon.exe. Dynamic range was set to 0.00-0.08, ring artifact correction was set to 16, and beam hardening was set to 30%. No smoothing was performed.

**Supplementary Figures**

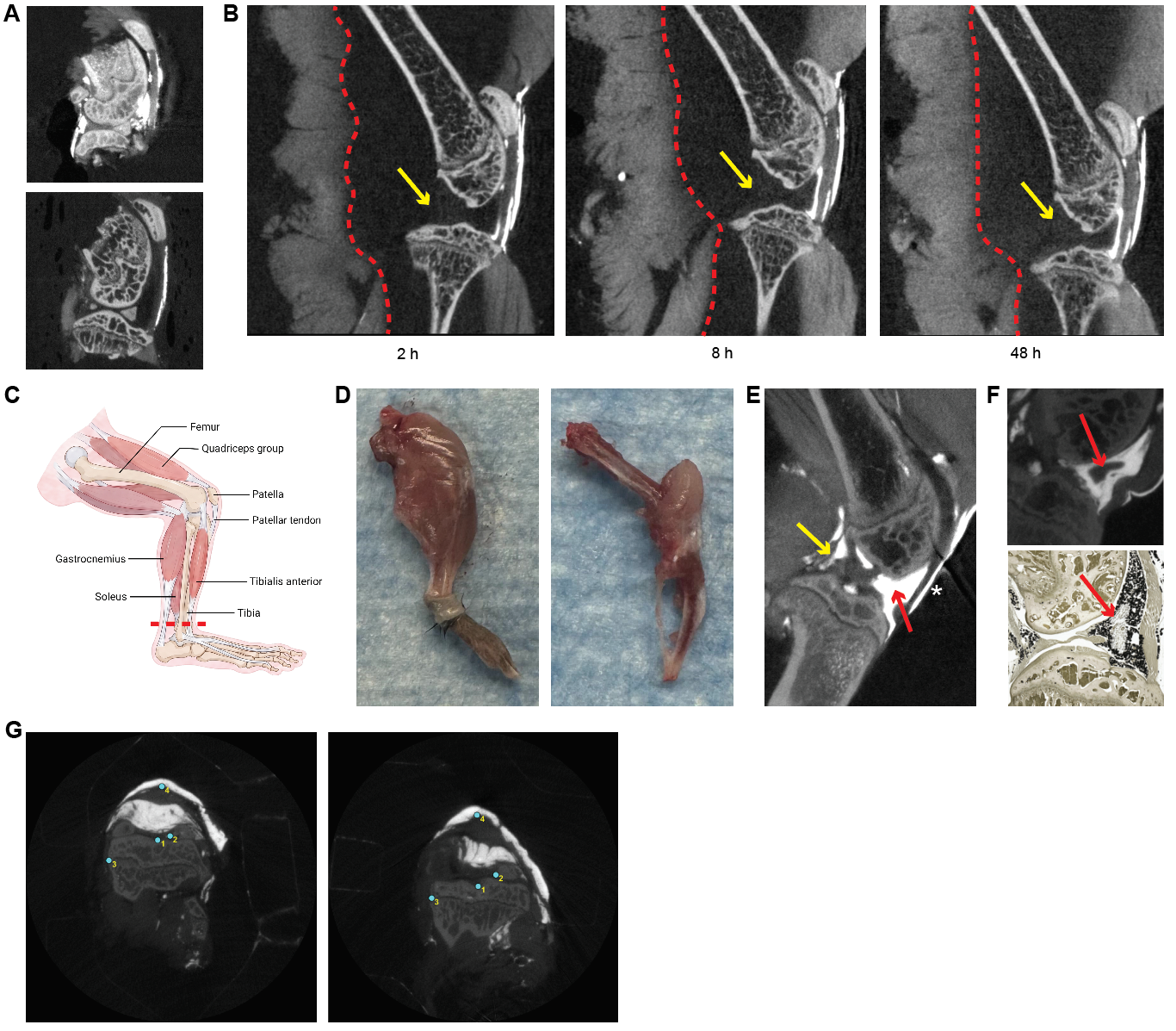

**Figure S1. Optimization of OsTet staining and μCT imaging.** (**A**) Representative μCT sagittal slices from pilot experiments demonstrating μCT detector saturation and beam hardening effects (top), and incomplete intra-articular adipose tissue staining (bottom). (**B**) Representative μCT sagittal slices of hindlimbs at increasing timepoints of OsTet staining. Dashed red lines show penetration depth of OsTet in muscle tissue. Yellow arrows show absent PFP staining. (**C**) Representative sagittal anatomical view of a mouse hindlimb, with red dashed line showing where tendons were cut then muscles removed. Created with BioRender.com. (**D**) Picture of mouse hindlimb prior to (left) and after (right) optimized dissection protocol. (**E**) Representative sagittal μCT slice of hindlimb following refinement of dissection procedure (removal of most muscle from limb). Yellow arrow points to positive staining of PFP; red arrow points to IFP; white asterisk is subcutaneous, extra-articular adipose tissue anterior to the joint. (**F**) Representative μCT sagittal slice of staining void in the IFP (top) and slice-matched OsTet histology showing unstained adipocytes (bottom). (**G**) Representative μCT axial slices with the four landmarks used in MATLAB for geometric registration between contralateral (left) and 28d ACLR (right) limbs.

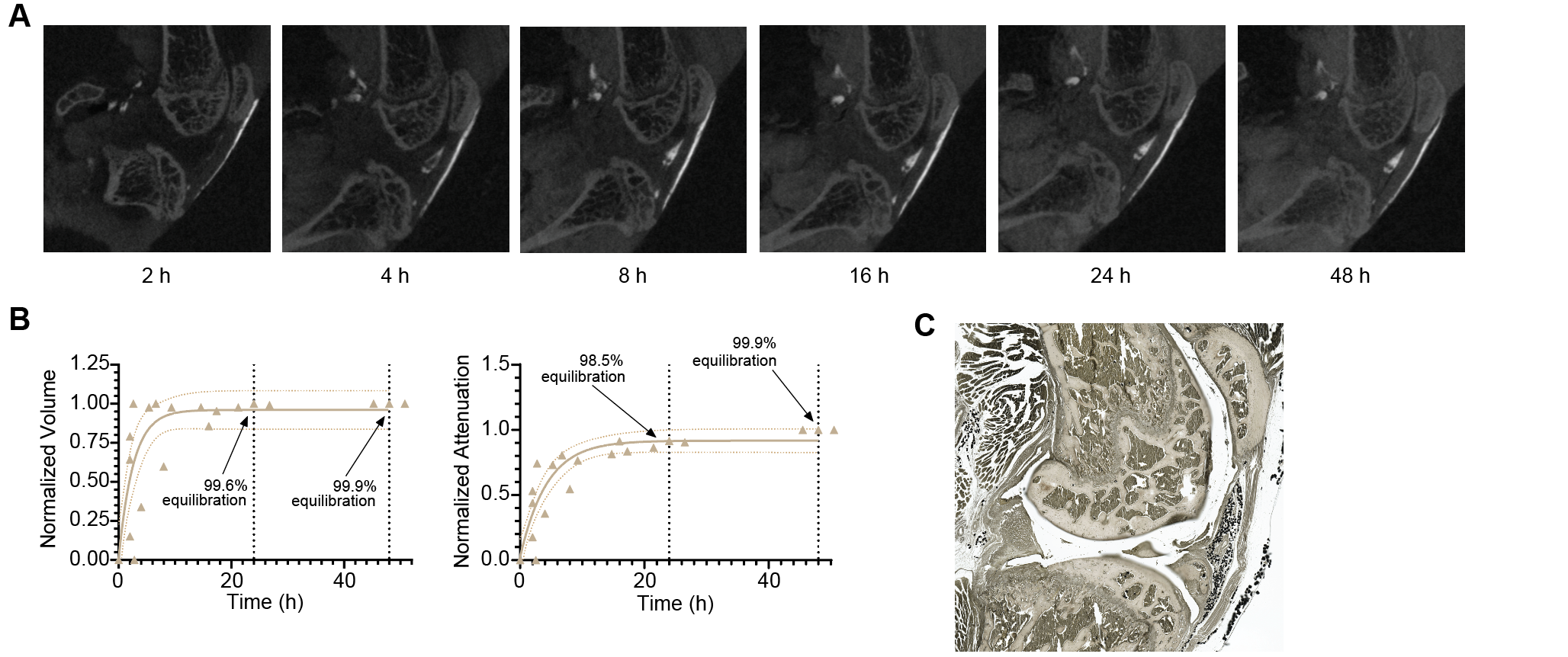

**Figure S2. OsTet staining kinetics in ACLR.** (**A**) Representative μCT slices from longitudinal scanning of 28d ACLR hindlimbs from 2-48 h. (**B**) Nonlinear regression modeling of normalized fat volume (left) and normalized attenuation (right) in 28d ACLR hindlimbs. Values are normalized to final volume or attenuation for each sample. Equilibration extents are shown at 24 h and 48 h (large vertical dotted lines) and 95% confidence interval are shown (small dotted lines). (**C**) Representative sagittal OsTet-stained histology section of 28d ACLR hindlimb.

**Supplementary Tables**

**Table S1.** Summary of nonlinear regression models assessing the relationship between equilibration and time for uninjured (contralateral) limbs.

| **Uninjured Hindlimb Regression Modeling** | **Metric** | **Volume-Based** | **Attenuation-Based** |
| --- | --- | --- | --- |
| Best-fit Values | Y_0_ | 0.02337 | 381.7 |
|  | Plateau | 0.8673 | 9205 |
|  | K | 0.1465 | 0.1203 |
|  | Tau (𝜏) | 6.825 | 8.316 |
|  | Half-time | 4.731 | 5.764 |
|  | Span | 0.8440 | 8823 |
| 95% Confidence Interval | Y_0_ | -0.07119 to 0.1146 | -542.1 to 1264 |
|  | Plateau | 0.7926 to 0.9539 | 8365 to 10214 |
|  | K | 0.1001 to 0.2089 | 0.08155 to 0.1736 |
|  | Tau (𝜏) | 4.786 to 9.990 | 5.760 to 12.26 |
|  | Half-time | 3.317 to 6.924 | 3.992 to 8.500 |
| Goodness of Fit | Degrees of Freedom | 18 | 18 |
|  | R^2^ | 0.9382 | 0.9433 |
|  | Sum of Squares | 0.1267 | 12533808 |
|  | Sy.x | 0.08390 | 834.5 |

**Table S2.** Summary of nonlinear regression models assessing the relationship between equilibration and time for 28d ACLR limbs.

| **28d ACLR Regression Modeling (normalized)** | **Metric** | **Volume-Based** | **Attenuation-Based** |
| --- | --- | --- | --- |
| Best-fit Values | Y_0_ | 0.005104 | 0.02333 |
|  | Plateau | 0.9606 | 0.9178 |
|  | K | 0.3856 | 0.2321 |
|  | Tau (𝜏) | 2.593 | 4.309 |
|  | Half-time | 1.798 | 2.987 |
|  | Span | 0.9555 | 0.8945 |
| 95% Confidence Interval | Y_0_ | -0.2223 to 0.2310 | -0.1227 to 0.1673 |
|  | Plateau | 0.8390 to 1.092 | 0.8269 to 1.022 |
|  | K | 0.1939 to 0.7776 | 0.1353 to 0.3733 |
|  | Tau (𝜏) | 1.286 to 5.158 | 2.679 to 7.393 |
|  | Half-time | 0.8914 to 3.575 | 1.857 to 5.124 |
| Goodness of Fit | Degrees of Freedom | 18 | 18 |
|  | R^2^ | 0.7828 | 0.8893 |
|  | Sum of Squares | 0.6360 | 0.2629 |
|  | Sy.x | 0.1880 | 0.1208 |
